# Improving cell-free glycoprotein synthesis by characterizing and enriching native membrane vesicles

**DOI:** 10.1101/2020.07.19.211201

**Authors:** Jasmine M. Hershewe, Katherine F. Warfel, Shaelyn M. Iyer, Justin A. Peruzzi, Claretta J. Sullivan, Eric W. Roth, Matthew P. DeLisa, Neha P. Kamat, Michael C. Jewett

**Author notes:** Authors contributed equally (Jasmine M. Hershewe, Katherine F. Warfel). To whom correspondence should be addressed Michael C. Jewett, **Email:**.

## Abstract

Cell-free gene expression (CFE) systems from crude cellular extracts have attracted much attention for accelerating the design of cellular function, on-demand biomanufacturing, portable diagnostics, and educational kits. Many essential biological processes that could endow CFE systems with desired functions, such as protein glycosylation, rely on the activity of membrane-bound components. However, without the use of synthetic membrane mimics, activating membrane-dependent functionality in bacterial CFE systems remains largely unstudied. Here, we address this gap by characterizing native, cell-derived membrane vesicles in *Escherichia coli*-based CFE extracts and describing methods to enrich vesicles with heterologous, membranebound machinery. We first use nanocharacterization techniques to show that lipid vesicles in CFE extracts are tens to hundreds of nanometers across, and on the order of ~3×10^12^ particles/mL. We then determine how extract processing methods, such as post-lysis centrifugation, can be used to modulate concentrations of membrane vesicles in CFE systems. By tuning these methods, we show that increasing the number of vesicle particles to ~7×10^12^ particles/mL can be used to increase concentrations of heterologous membrane protein cargo expressed prior to lysis. Finally, we apply our methods to enrich membrane-bound oligosaccharyltransferases and lipid-linked oligosaccharides for improving *N*-linked and *O*-linked glycoprotein synthesis. We anticipate that our findings will facilitate *in vitro* gene expression systems that require membrane-dependent activities and open new opportunities in glycoengineering.

## Introduction

Lipid membranes play pivotal roles in biological functions across all domains of life, with ~20-30% of genes encoding for membrane proteins and many essential processes taking place on and across membranes (1,2). For example, membranes are required for molecular transport, immunological defense, energy regeneration, and post-translational protein modification. Despite the absence of intact cellular membranes, crude extracts of organisms have proven useful in a variety of *in vitro* studies by reconstituting some of these biological phenomena. This is possible due to the presence of membrane structures which form upon fragmentation and rearrangement of cell membranes during cell lysis and extract preparation. In eukaryotic-derived cell-free gene expression (CFE) systems, endoplasmic reticulum (ER)-derived microsomes enhance functionality, enabling the synthesis of membrane proteins, proteins with disulfide bonds, and protein glycosylation (3–7). Eukaryotic ER microsomes have been routinely characterized for quality control with fluorescence microscopy due to their micron-scale size, enabling the development of diverse systems that leverage microsomes (8–10). Analogously, in *E. coli-derived* CFE systems, inverted membrane vesicles harboring electron transport chain machinery activate oxidative phosphorylation and ATP regeneration (11, 12). While vesicles in typical *E. coli* extracts have been analyzed using biochemical methods, sucrose fractionation, and phospholipid quantitation, the use of methods to count and characterize intact, cell-derived vesicles have not yet been pursued (13–15). This is due, in part, to the fact that vesicles in *E. coli* extracts are on the nanoscale and therefore require higher-resolution techniques than fluorescence microscopy for characterization. While exogenous membranes such as nanodiscs, synthetic phospholipid structures, purified microsomes, and purified vesicles have enabled membrane biology in CFE systems (16–21), using native membranes from the host simplifies processing and is an appealing alternative. Therefore, characterization workflows for analyzing native vesicles are a foundational step for enabling applications of membrane-bound biology in *E. coli-based* CFE.

A technical renaissance has recently transformed CFE systems from a molecular biology technique to a widely applicable bioproduction and prototyping platform, with *E. coli* systems at the forefront (4, 22–27). A body of work dedicated to optimization of extract preparation and reaction conditions has simplified, expedited, and improved the cost and performance of *E. coli* CFE systems (22, 28, 29). Optimized *E. coli-based* CFE reactions: (i) quickly synthesize grams of protein per liter in batch reactions (30–32), (ii) are scalable from the nL to 100 L scale (33, 34), and (iii) can be freeze-dried for months of shelf-stability and distribution to the point of care (6, 22, 29, 35–40). Freeze-dried CFE systems are poised to make disruptive impacts in biotechnology, having already been leveraged for point-of-use biosensing (41–46), therapeutic and vaccine production (37, 38, 47), and educational kits (22, 48–50).

In a growing number of contexts, CFE extracts have been tailored to new applications by pre-enriching soluble, heterologous components *in vivo* prior to cell lysis, avoiding the need for purification. Examples include incorporation of site-specific non-canonical amino acids into proteins (31, 51, 52), biosensing of analytes (45, 53, 54), and assembly of metabolic pathways for production of valuable small molecules (26, 55–57). Yet, the design of membrane-incorporated components to enhance CFE systems has remained largely unstudied. Enriching membranebound components in CFE systems would enable compelling applications. For example, protein glycosylation, which is mediated by membrane-bound components, is a key consideration in cell-free biomanufacturing of protein therapeutics and conjugate vaccines. We recently described cell-free glycoprotein synthesis (CFGpS), a platform for one-pot biomanufacturing of defined glycoproteins in extracts enriched with heterologous, membrane-bound glycosylation machinery (47). To date, CFGpS has been used to produce model glycoproteins, human glycoproteins, and conjugate vaccines (38, 47, 58–60). Importantly, CFGpS reactions can be freeze-dried for shelf-stability and rehydrated at the point of care to make effective vaccines (38). Because CFGpS activity relies on membrane-bound glycosylation components, methods to characterize and quality-control the membrane-bound components in extracts is paramount for moving the technology forward.

In this work, we characterize size distributions and concentrations of native membrane vesicles in extracts, providing a benchmark for analysis and engineering of CFE systems. We investigate the impacts of upstream extract processing steps on vesicle profiles, revealing simple handles to modulate vesicle concentration in extracts. We use native membrane vesicles to enrich a variety of heterologous, membrane-bound proteins and substrates in extracts without the use of synthetically-derived membranes. Finally, we apply our findings to improve glycoprotein yields in our existing asparagine-linked (*N-*linked) CFGpS system and a new membrane-dependent CFGpS system based on serine/threonine-linked (*O*-linked) glycosylation. The implications of our work extend beyond glycosylation and are applicable to engineering new CFE systems with membrane-associated activities.

## Results

In this study, we aimed to characterize membrane vesicles in *E. coli-based* CFE systems that form upon fragmentation of cell membranes during cell lysis. Then, we wished to use this knowledge to control enrichment of membrane-bound components for enhancing defined function. To achieve these goals, we (i) use nanocharacterization techniques to determine the sizes and quantities of membrane vesicles in *E. coli* extracts; (ii) determine how extract processing can control the enrichment of vesicles in extracts; (iii) enrich several heterologous, membrane-bound components in extracts via vesicles; and (iv) demonstrate that enrichment of membrane-bound components improves cell-free glycoprotein synthesis systems for *N-* and *O*-linked glycosylation.

### Characterization of membrane vesicles in CFE extracts

Initially, we used several nanocharacterization techniques to analyze the size of vesicles and visualize these particles in CFE extracts. Dynamic light scattering (DLS) analysis of crude extract revealed two major peaks: one narrower peak with an intensity maximum at ~20 nm, and a broader peak at ~100-200 nm (**Fig. 1A**). The 20 nm peak represents small cell-derived particles, including assembled 20 nm *E. coli* ribosomes (61), which we confirmed to be active in our extracts (**SI Appendix, Fig. S1**). We hypothesized that particles measured in the ~100-200 nm peak were vesicles. An illustration of particles detected in extract is shown in **Fig. 1B**. To directly analyze membrane vesicles without ribosomes and other cellular particles, we identified and purified membranous particles via size exclusion chromatography (SEC) (62–64) (**SI Appendix, Fig. S2A**). DLS analysis of purified membrane vesicles revealed an intensity particle size distribution (PSD) that directly overlapped with the proposed vesicle peak from our DLS traces of crude extracts. (**Fig. 1A**). Nanoparticle Tracking Analysis (NTA), an orthogonal method for sizing and quantitating nanoparticles in solution, revealed an average purified vesicle diameter of 118.5±0.7 nm, corroborating the size range measured with DLS (**SI Appendix, Fig. S2B**). The zeta potential of purified vesicles was −14.5±-1.0 mV, indicating a negative particle surface charge consistent with phospholipid vesicles (**SI Appendix, Fig. S2C**). Cryo-electron microscopy (cryo-EM) of extracts showed small (≤20 nm) particles and other larger, circular particles consistent with vesicle morphology (**Fig. 1C**). Cryo-EM micrographs of extracts revealed vesicles between ~40 nm and ~150 nm in size, and morphologies were consistent pre- and post-SEC purification (**Fig. 1C-1D**). Comparisons between measurements reveal that DLS, a bulk, in-solution measurement, over-estimates vesicle diameter. DLS, however, is a useful tool for quickly characterizing crude extract particle profiles because it can detect particles <50 nm (including ribosomes) that are smaller than vesicles and are below the size limit of detection of NTA. Together, these results show particle profiles of crude extracts and reveal that vesicles are polydisperse, are on the order of tens to hundreds of nm across, and are relatively low in concentration compared with ribosomes and other small complexes.

**Figure 1.**
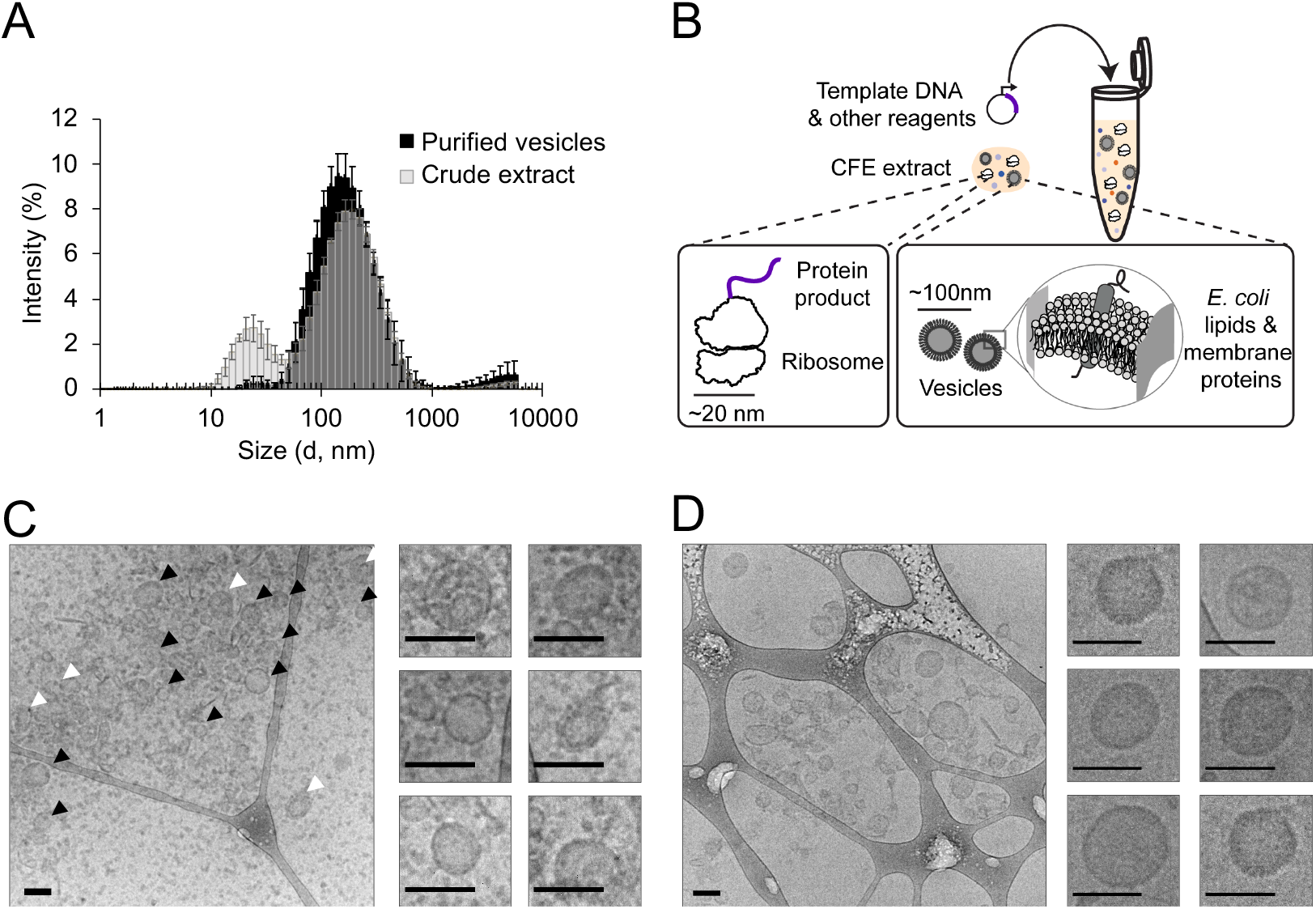
Characterization of membrane vesicles in crude CFE extracts. **(A)** DLS analysis of crude extracts and SEC purified vesicles. Crude extract trace is translucent to show overlap. Error bars represent the standard deviation within triplicate analysis of three independently prepared extracts. For purified vesicles, error bars represent the standard deviation of triplicate analysis of the most concentrated vesicle elution fractions. **(B)** Illustration of particles detected in crude CFE extracts. **(C)** Cryo-EM micrographs of crude extracts. Black arrows indicate vesicles with apparent unilamellar morphology. White arrows indicate nested or multilamellar morphologies. Cropped images indicate representative vesicles. Scale bars are 100 nm. **(D)** Cryo-EM micrographs of SEC purified vesicles. Cropped images indicate representative purified vesicles. Scale bars are 100 nm.

### Extract processing impacts vesicle size distributions and concentrations

To understand how to control membrane vesicles in extracts, we next sought to study how protocols to process extracts impacted vesicle properties. Specifically, we studied cell lysis and extract centrifugation because cell membranes are ruptured during lysis, and centrifugation dictates particle separation. We lysed cells using standard sonication (constant input energy per volume of cell suspension) or homogenization protocols (~20,000 psig) (28, 47), then subjected lysates to a traditional 30,000 x *g* centrifugation protocol (termed ‘S30 prep’), or a lower g-force protocol where the maximum centrifugation speed was 12,000 x *g* (termed ‘S12 prep’) (**Fig. 2A**) (28, 29). These combinations of lysis and centrifugation protocols resulted in four distinct extract conditions, all of which were active for protein synthesis in standard CFE reaction conditions (**SI Appendix, Fig. S3A**). The combination of a standard homogenization and S30 prep represents our base case because extracts used in our previously described one-pot cell-free glycoprotein synthesis platform were prepared with these conditions, as well as the extracts used in **Fig. 1**.

**Figure 2.**
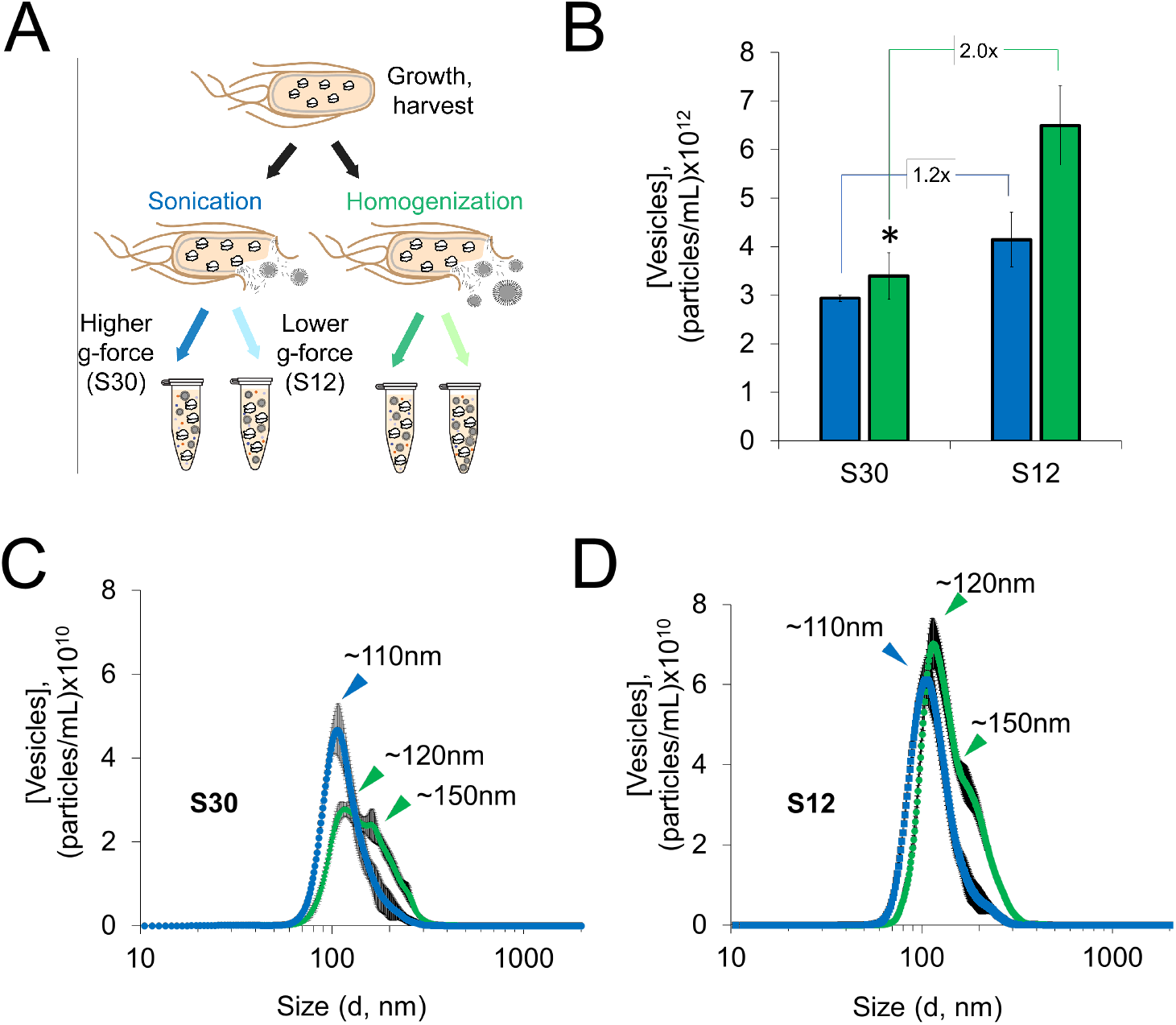
Extract processing impacts vesicle size distributions and concentrations. **(A)** Illustration of extract processing conditions. Extracts were prepared in triplicate for each condition shown. **(B)** Nanoparticle Tracking Analysis (NTA) concentration analysis of vesicles in sonicated (blue) and homogenized (green) extracts. Asterisk indicates base case conditions for extract preparation. For NTA analyses, error bars represent the standard deviation of measurements of three independently prepared extracts. **(C)** NTA particle size distribution (PSD) of vesicles in sonicated (blue) and homogenized (green) S30 extracts. **(D)** NTA PSD of sonicated (blue) and homogenized (green) S12 extracts.

Of the conditions tested, the centrifugation protocol had the most impact on vesicle concentrations. We observed significantly higher numbers of vesicles in S12 extracts for both lysis methods. Specifically, we observed 1.2- and 2.0-fold enrichments of vesicles in sonicated and homogenized S12 extracts, respectively (**Fig. 2B**). Homogenized S12 extracts contained the highest concentration of vesicles with 6.5±0.8×10^12^ particles/mL (as compared to 3.4±05×10^12^ particles in the base case), making it the most promising condition for enriching vesicles.

While centrifugation impacted vesicle concentration, lysis method impacted vesicle size. Sonicated extracts contained smaller vesicles with narrower size distributions than homogenized extracts, regardless of centrifugation protocol. Our observations that lysis method impacts vesicle size is consistent with studies showing that varying experimental parameters to disperse phospholipids (or amphiphiles in general) impacts vesicle sizes (65). PSDs of sonicated extracts reached single maxima at ~110 nm, with average particle diameters of ~130nm; homogenized extracts had higher average particle diameters of ~160 nm, displaying distinct peaks at ~120 nm, and considerable shoulder peaks at ~150 nm (**Fig. 2C–2D, SI Appendix, Fig S4A**). Homogenized PSDs may indicate the presence of multiple, discrete, vesicle populations (**Fig 2C–2D)**. DLS measurements confirmed the observation that sonicated extracts contained relatively smaller, less polydisperse vesicles than homogenized extracts (**SI Appendix, Figs. S4B-S4C**). Notably, direct vesicle analysis in extracts enabled us to gauge the impacts of extract processing in ways that have not been previously accessible and provides benchmarks for intact vesicle concentrations in extracts.

### Heterologous membrane-bound cargo can be controllably enriched via membrane vesicles

With a better understanding of the characteristics and concentrations of native vesicles, we sought to enrich extracts with vesicles containing heterologous cargo derived from the periplasmic membrane of *E. coli.* Since S12 extracts contain higher concentrations of vesicles than S30 extracts, we hypothesized that S12 extracts would also contain higher concentrations of associated heterologous cargo. The highest dynamic range of vesicle concentration between S12 and S30 preps was observed with homogenization, so we proceeded with homogenization for enrichment experiments (**Fig. 2B**). We overexpressed six membrane-bound proteins of various sizes, transmembrane topologies, biological functions, and taxonomical origins to test for enrichment (**SI Appendix, Table S1**). The proteins selected for enrichment encompass classes of proteins that could enable new functionalities in CFE, including glycosylation enzymes (PglB, PglO, STT3) and signal transduction/sensing proteins (NarX, PR, CB1). We expressed each membrane protein *in vivo* with a C-terminal FLAG tag, prepared S30 and S12 extracts, then analyzed concentrations of the overexpressed membrane protein using quantitative Western blotting. We observed 2-fold membrane protein enrichment in S12 over S30 (S12/S30) extracts for all proteins other than PR, for which we observed 4-fold enrichment (**Fig 3A–3B**). As a control, when sfGFP with no transmembrane helices was expressed *in vivo,* we did not observe significant S12/S30 enrichment (**Fig. 3C**). Full blots for **Fig. 3A–3C** are shown in **SI Appendix, Fig. S5**. Notably, enrichment values obtained via blotting correspond closely with the 2-fold vesicle enrichment observed via NTA in homogenized S12 and S30 extracts with no overexpression (**Fig. 2B**). All extracts with pre-enriched membrane proteins displayed protein synthesis activity (**SI Appendix, Fig. S3B**).

**Figure 3.**
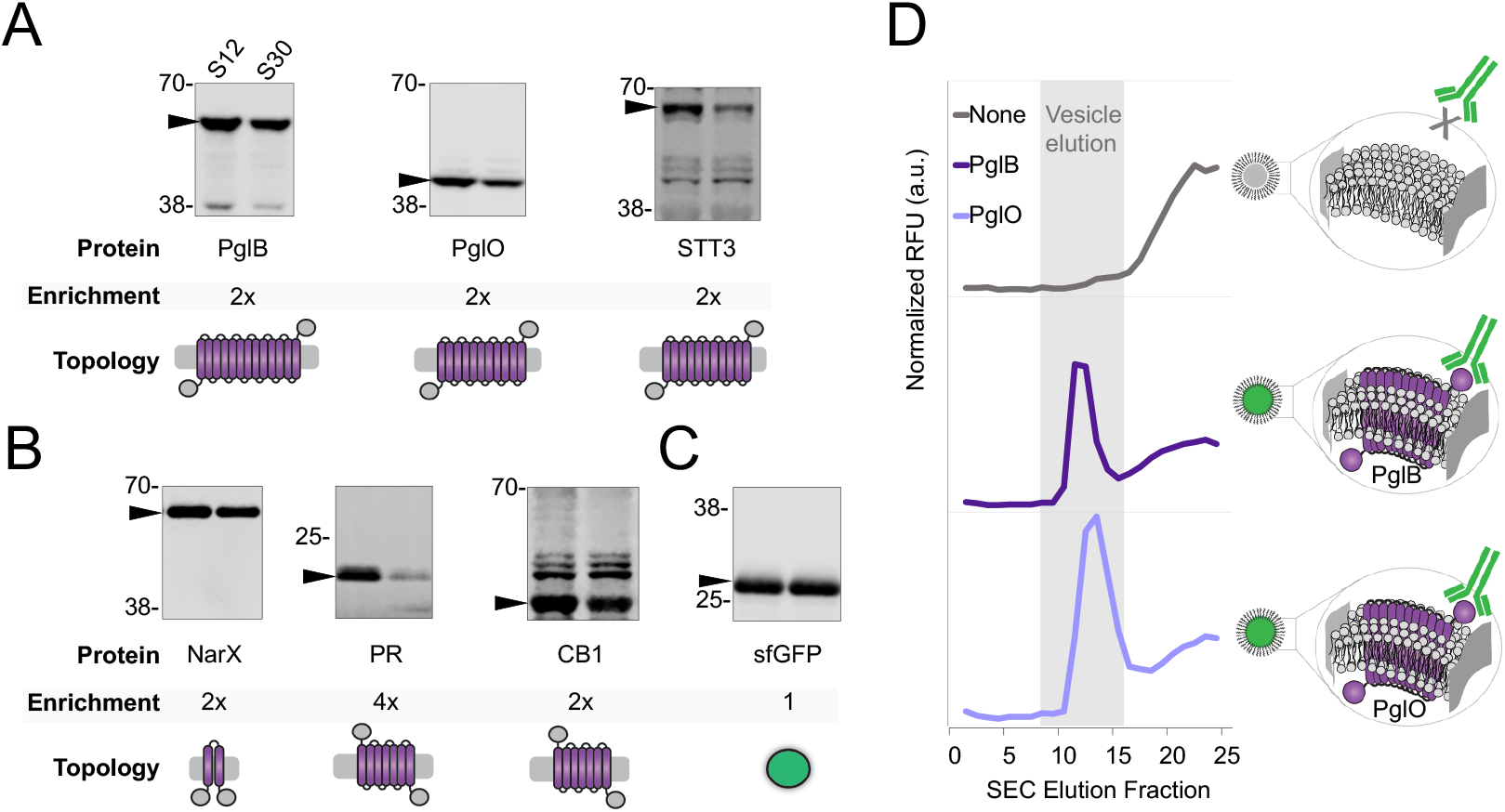
Heterologous membrane-bound cargo can be controllably enriched via membrane vesicles. Enrichment of heterologous membrane proteins in S12 and S30 extracts was quantitated using α-FLAG Western blots against the heterologous proteins for **(A)** glycosylation enzymes and **(B)** signal transduction proteins. **(C)** Cytosolic sfGFP control with no transmembrane helices. On each Western blot, left lanes are S12 extracts and right lanes are S30 extracts. Black arrow indicates the membrane protein of interest. Molecular weight (kDa) from protein ladder standards are indicated to the left of each blot. Protein names and enrichment ratio of bands (S12/S30) are shown directly below each blot. All blots are representative of 3 independent experiments. Cartoons depict the transmembrane topology for each protein. See **Table S1** for taxonomical origin, transmembrane topology, functions(s), theoretical size, and UniProt ID. **(D)** Fluorescence chromatograms of SEC analysis of extracts probed with a fluorescent α-FLAG antibody. Strains used to prepare extracts were enriched with no membrane protein (gray trace), PglB (dark purple trace), or PglO (light purple trace). Characteristic vesicle elution fraction from 3 independent experiments is highlighted in gray.

We next confirmed that PglB and PglO, key enzymes for glycosylation, were associated with membrane vesicles, as opposed to free in solution (**Fig. 3D**). Extracts with pre-enriched PglB or PglO were probed with a green fluorescent α-FLAG antibody, then analyzed via SEC. Fluorescence chromatograms are shown in **Fig. 3D**, with the characteristic vesicle elution fraction highlighted in gray (**SI Appendix, Fig. S2A**). The characteristic vesicle elution peak corresponded with green fluorescence for extracts containing PglB or PglO and no corresponding peak was observed in an extract with no overexpressed membrane protein (**Fig. 3D**). Our results show that heterologous, periplasmic membrane cargo can be pre-enriched in extract and tuned via vesicles.

### Increasing vesicle concentrations improves cell-free glycoprotein synthesis (CFGpS) for N- and O-linked glycosylation systems

We next set out to exploit our ability to enrich vesicles harboring heterologous cargo in an application. We focused on protein glycosylation, because glycosylation plays critical roles in cellular function, human health, and biotechnology. As a model, we sought to increase glycoprotein yields in a previously reported CFGpS platform (47) by charging reactions with S12 extracts containing higher concentrations of membrane-bound glycosylation machinery. We prepared S30 and S12 extracts from strains overexpressing the model *N-*linked glycosylation pathway from *Campylobacter jejuni,* which consists of the membrane-bound oligosaccharyltransferase (OST) PglB that catalyzes glycosylation, and a lipid-linked oligosaccharide (LLO) donor of the form: GalNAc-α1,4-GalNAc-α1,4-(Glcβ1,3)-GalNAc-α1,4-GalNAc-α1,4-GalNAc-α1,3-Bac (where Bac is 2,4-diacetamido-2,4,6-trideoxyglucopyranose) from an undecaprenylpyrophosphate-linked donor (66). NTA and Western blot analysis of CFGpS extracts revealed 2.5-fold S12/S30 enrichment of vesicles and a corresponding 2-fold S12/S30 enrichment of PglB (**SI Appendix, Fig. S6**). Fluorescence staining and SEC analysis confirmed the presence of LLO and PglB in vesicles (**SI Appendix, Fig. S7A**).

To assess the impact of enriched vesicles on cell-free glycoprotein synthesis, we carried out reactions in two phases (**Fig. 4A, Inset**) (38). First, cell-free protein synthesis (CFPS) of the acceptor protein was run for a defined time, termed ‘CFPS time’. At the CFPS time, reactions were spiked with MnCl_2_, quenching CFPS and initiating glycosylation by providing the OST with its Mn^2+^ cofactor. CFGpS reactions charged with S30 or S12 extracts were run for CFPS times of 2, 10, 20, 30, and 60 minutes using a His-tagged sfGFP_DQNAT_ acceptor protein, where DQNAT is a permissible PglB sequon. Coding sequences of all acceptor proteins used are presented in **SI Appendix, Table S2**. Endpoint glycoprotein yields were quantified using total acceptor protein fluorescence and % glycosylation determined by Western blotting (**Fig. 4A, SI Appendix, Fig. S8A-Fig.S8D)**. At longer CFPS times we observed that S12 extracts produced significantly more glycoprotein than S30 extracts. Because total acceptor protein concentrations for S30 and S12 reactions were similar for each CFPS time (**SI Appendix, Figure S8E)**, increased glycoprotein yield in S12 extracts is due to higher glycosylation activity and not higher CFE yields. Specifically, at 20, 30, and 60-minute CFPS times, we observed 66%, 85%, and 90% increases in glycoprotein yield in the S12 reactions, respectively. At the 60-minute CFPS time, S12 reactions yielded 117.2±9.9 μg/mL in batch, synthesizing glycoprotein titers on the order of hundreds μg/mL for the first time to our knowledge. (**Figure 4A**). S12 reactions also had significantly higher terminal % glycosylation, or percent of CFPS-derived acceptor protein that is glycosylated at the end of a 16hour glycosylation reaction. This was true for all CFPS times tested (**SI Appendix, Figure S8F**).

**Figure 4.**
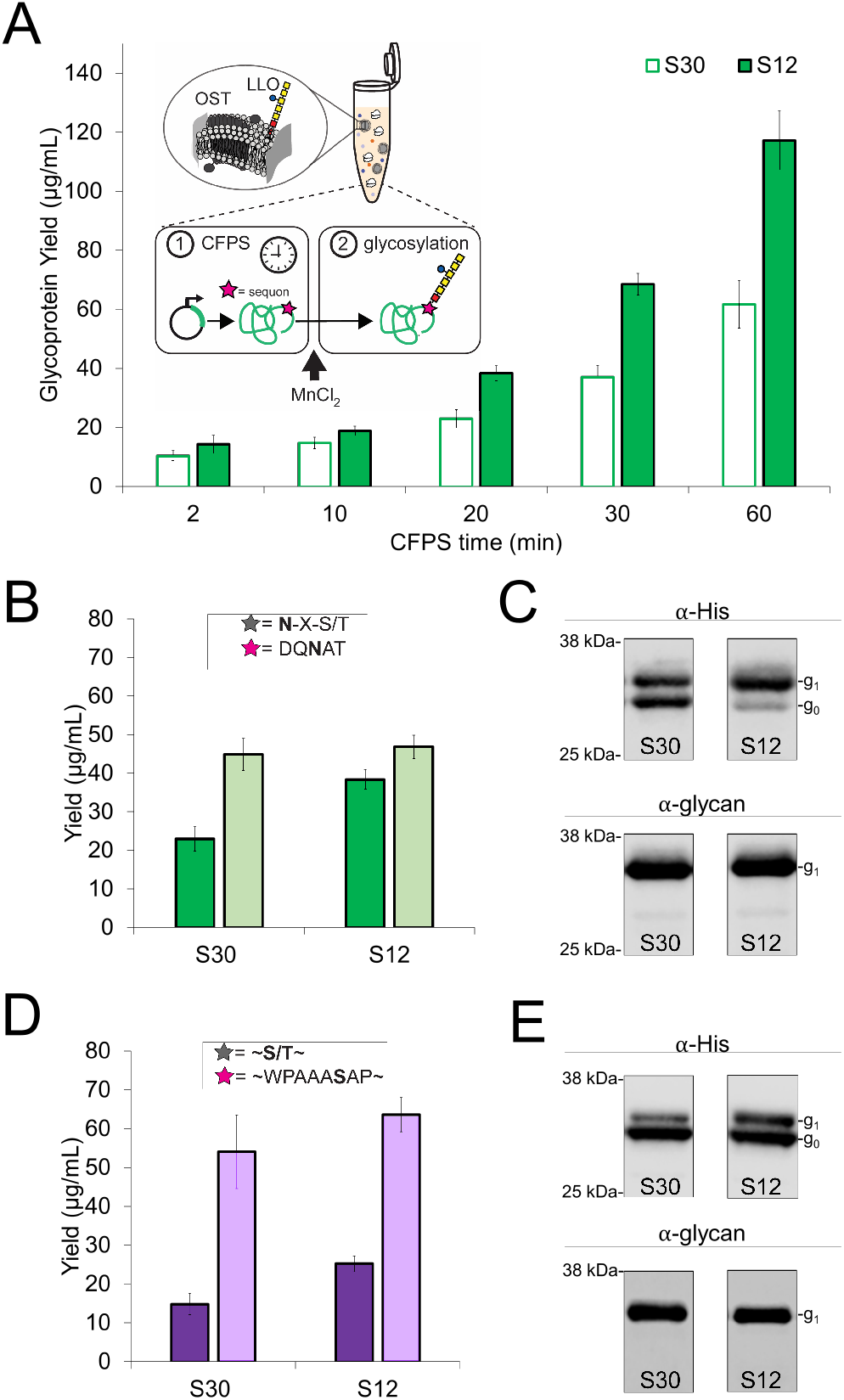
Increasing vesicle concentrations improves cell-free glycoprotein synthesis (CFGpS) for *N*- and *O*-linked glycosylation systems. **(A)** Glycoprotein yields of CFGpS reactions charged with S12 or S30 extracts enriched with PglB and *C. jejuni* LLO. Error bars represent standard deviation of 3 independent CFGpS reactions, each run with an independent extract. **(Inset)** Schematic of 2-phase CFGpS reactions. **(B)** Glycosylated (dark green) and total (light green) protein yields of *N*-linked CFGpS reactions with 20 minute CFPS times. Error bars represent standard deviation of 3 independent reactions. **(Inset)** Sequon preferences for general *N*-OST mediated (gray star) and PglB (pink star) mediated glycosylation. Glycosylated residue is bolded. **(C)** Western blots of acceptor proteins from representative reactions in **(B)**, where g_o_ denotes aglycosylated acceptor protein and gi indicates glycoprotein. **(D)** Glycosylated (dark purple) and total (light purple) protein yields from *O*-linked CFGpS reactions with 20 minute CFPS times. **(Inset)** Sequon preferences for general *O*-OST (gray star) and PglO (pink star) mediated glycosylation. Glycosylated residue is bolded. **(E)** Western blots of acceptor proteins from representative reactions in **(D)**. All gels are representative of 3 independent CFGpS reactions.

For example, for reactions with 20-minute CFPS times, we observed an increase from 51% glycosylation for S30 reactions to 82% glycosylation for S12 reactions (**Fig. 4B**). α-His (showing glycosylated and aglycosylated acceptor protein) and α-glycan (against the *C. jejuni* glycan) Western blots of representative reactions are shown in **Fig. 4C.** Taken together, these results indicate that the presence of higher vesicle concentrations in S12 extracts has a measurable effect on CFGpS, improving glycoprotein yields and endpoint % glycosylation.

With a long-term interest in synthesizing diverse glycoproteins in cell-free systems, we next ported an *O*-linked glycosylation system, known to have broad glycan specificity, into the CFGpS platform (60, 67, 68). We selected the *O*-OST PglO from *Neisseria gonorrhoeae* which accepts the *C. jejuni* heptasaccharide LLO as a donor but differs from PglB in acceptor sequence preferences (69). **Fig. 4B** and **Fig. 4D** insets show sequons for PglB and PglO, respectively. For PglO, we used an sfGFP-fusion acceptor protein containing a recently determined 8 amino acid minimum optimal *O*-linked recognition site (termed ‘MOOR’) (69). We confirmed residue-specific *O-*linked glycosylation and enrichment of PglO and LLO in vesicles (**SI Appendix, Fig. S9, Fig. S7B**). As in PglB-mediated CFGpS, we observed increased endpoint glycoprotein yield and % glycosylation in reactions charged with S12 extracts. Specifically, reactions with CFPS times of 20 minutes resulted in a 68% increase in glycoprotein yield and an increase from 27% to 40% glycosylation in reactions with S12 extracts compared to those containing S30 extracts (**Fig. 4D**, **SI Appendix, Fig. S10**). Corresponding blots are shown in **Fig. 4E** and **SI Appendix, Fig. S10A-S10B.** Taken together, these results indicate that improvements to glycosylation in S12 extracts translate from the *N-*linked glycosylation system to the *O*-linked glycosylation system.

## Discussion

In this work, we set out to benchmark, understand, and quality-control protein-enriched vesicles in CFE extracts for expanding and enhancing functionality. We showed that upstream extract processing can be used to tune concentrations of vesicles and associated cargo from the periplasm. Then, we applied this knowledge to improve cell-free glycoprotein synthesis. Our results have several key features.

First, the nanocharacterization tools used here allowed us to quantify intact vesicle numbers and sizes in CFE extracts. This is important because this knowledge informed design rules for enhancing vesicle concentrations and functionality from their associated protein cargo in cell-free systems. Notably, the effective vesicle surface area calculated from NTA measurements (~0.3 m^2^ membrane/mL extract), is consistent with values calculated from phospholipid concentrations in similar extracts (15).

Second, our results offer insights into field-wide observations and limitations of *E. coli-* based CFE systems. For example, it is well-documented that lysis protocols can profoundly impact CFE productivity (70). Our findings show that lysis methods impact size distributions of vesicles generated during this step which affect the membrane environment of the machinery necessary for oxidative phosphorylation and ATP regeneration. Since vesicles are key for activating cost-effective energy metabolism from oxidative phosphorylation in CFE, routine vesicle characterization could become a vital quality-control check, leading to improved reproducibility in and between labs (71). Our results also offer insight into why, despite the presence of vesicles in the *E. coli* CFE system, CFE-derived membrane proteins cannot be synthesized via insertion into native vesicles without additional vesicle supplementation (12, 19, 21). With ~6 nM of intact vesicles in CFE reactions (estimated from NTA measurements), the concentration of vesicles is orders of magnitude lower than typical protein titers produced in our CFE extracts (~30 μM of reporter protein or higher).

Third, our work opens the door to engineering systems that rely on enriched membrane-bound components. We show that membrane-bound proteins and lipid-linked oligosaccharides expressed *in vivo* in the periplasm can be enriched in vesicles, indicating that a population of vesicles is derived from the periplasmic membrane (72, 73). Based on proteomic identification of both outer- and inner-membrane proteins in typical S30 extracts, and the presence of endotoxin in *E. coli* extracts (74), we hypothesize that multiple orientations and compositions (e.g., in/everted and inner/outer membrane) of vesicles may exist. Parsing out these details will require future characterization. Additionally, our workflow easily interfaces with additives that change or alter membrane properties (e.g., composition, size, fluidity, curvature) that could be added to the lysis buffer to tune the biophysical features of vesicles to further improve recombinant enzyme activity (19). And, while we focus entirely on *E. coli*-based systems here, the reported characterization methods could, in principle, be extended to further optimize insect and CHO-based CFE systems that rely on ER-derived microsomes to perform glycosylation, embed nascent membrane proteins, and perform other membrane-dependent functions.

Looking forward, we anticipate that our work will accelerate efforts to manufacture proteins that require membrane-dependent modifications, such as glycoproteins. For example, the approach described enables *N*-linked glycoprotein synthesis yields of >100 μg/mL, which increases accessibility for on-demand vaccine production in resource-limited settings. Further, the S12 prep does not require a high-speed centrifuge and is less time-intensive than the base case, simplifying the the CFGpS platform. Taken together, our results pave the way for efficient, accessible CFE systems that require membrane-bound activities for expanding system functionality and enabling a variety of synthetic biology applications.

## Materials and Methods

### Extract preparation

The chassis strain used for all extracts was CLM24 (47). Source strains were grown in 1 L of 2×YTPG media at 37 °C with agitation. Cells were grown to OD 3, then harvested by centrifugation (5000 × *g*, 4 °C, 15 min). For overexpression of proteins *in vivo,* CLM24 source strains were grown at 37 °C in 2xYTPG with the appropriate antibiotic(s), listed in **Table S3.** Cells were induced with 0.02% (w/v%) L-arabinose at OD 0.6-0.8, shifted to 30 °C, and harvested at OD 3. All subsequent steps were carried out at 4 °C and on ice unless otherwise stated. Pelleted cells were washed 3 times in S30 buffer (10 mM Tris acetate pH 8.2, 14 mM magnesium acetate, 60 mM potassium acetate). After the last wash, cells were pelleted at 7000 *× g* for 10 min, flash-frozen and stored at −80 °C. After growth and harvest, cells were thawed and resuspended to homogeneity in 1 mL of S30 buffer per gram of wet cell mass. For homogenization, cells were disrupted using an Avestin EmulsiFlex-B15 high-pressure homogenizer at 20,000-25,000 psig with a single pass (Avestin, Inc. Ottawa, ON, Canada). For sonication, input energy was calculated using an empirical correlation as described previously (28). Cells were sonicated on ice using a Q125 Sonicator (Qsonica, Newtown, CT) with a 3.175 mm diameter probe at a frequency of 20 kHz and 50% of amplitude. Energy was delivered to cells in pulses of 45 s followed by 59 s off until the target energy was delivered. Cells were lysed and clarified in triplicate. For S30 prep, lysed cells were centrifuged twice at 30,000 *× g* for 30 min; supernatant was transferred to a fresh tube for each spin. Supernatants were incubated with 250 rpm shaking at 37 °C for 60 min for runoff reactions. Following runoff, lysates were centrifuged at 15,000 × *g* for 15 min. Supernatants were collected, aliquoted, flash-frozen, and stored at −80 °C for further use. For S12 prep, lysed cells were centrifuged once at 12,000 *x g* for 10 min; supernatants were collected and subjected to runoff reactions as described above. Following runoff, lysates were centrifuged at 10,000 *× g* for 10 min at 4 °C. Supernatants were collected, aliquoted, flash-frozen in liquid nitrogen, and stored at −80 °C.

### Dynamic light scattering (DLS) and nanoparticle tracking analysis (NTA) measurements

DLS measurements were performed on a Zetasizer Nano ZS (Malvern Instruments Ltd., UK) with a measurement angle of 173° in disposable cuvettes (Malvern Instruments Ltd., UK ZEN0040). All measurements were collected in triplicate for 13 scans per measurement. Refractive index and viscosity were obtained from the instrument’s parameter library. The instrument’s ‘General Purpose’ setting was used to calculate intensity and number particle size distributions. For DLS of crude extracts, extracts were diluted 1:10 with 0.1 μm filtered PBS before analysis. For purified vesicle samples, elutions were analyzed directly without dilution.

NTA measurements were performed on a Nanosight NS300 using a 642 nm red laser (Malvern Instruments Ltd., UK). Samples were diluted to manufacturer-recommended particle concentrations in sterile PBS until a linear trend between dilution factor and concentration measured was found. Samples were flowed into the cell, and the instrument was focused according to manufacturer recommendations. Measurements were collected at room temperature, using a 1 mL syringe and a syringe pump infusion rate of 30 (arbitrary units). Data for each sample was collected in 5 separate 1 min videos, under continuous flow conditions. Mean particle diameters and particle concentrations were obtained from aggregate Nanosight experiment reports of each run, then averaged across triplicates and corrected for dilution factor.

### Transmission electron microscopy

For cryo-TEM measurement, 200 mesh Cu grids with a lacey carbon membrane (EMS Cat. # LC200-CU) were placed in a Pelco easiGlow glow discharger (Ted Pella Inc., Redding, CA, USA) and an atmosphere plasma was introduced on the surface of the grids for 30 seconds with a current of 15 mA at a pressure of 0.24 mbar. This treatment creates a negative charge on the carbon membrane, allowing for aqueous liquid samples to spread evenly over of the grid. 4 μL of sample was pipetted onto the grid and blotted for 5 seconds with a blot offset of +0.5 mm, followed by immediate plunging into liquid ethane within a FEI Vitrobot Mark III plunge freezing instrument (Thermo Fisher Scientific, Waltham, MA, USA). Grids were then transferred to liquid nitrogen for storage. The plunge-frozen grids were kept vitreous at –172 °C in a Gatan Cryo Transfer Holder model 626.6 (Gatan Inc., Pleasanton, CA, USA) while viewing in a JEOL JEM1230 LaB6 emission TEM (JEOL USA, Inc., Peabody, MA,) at 120 keV. Image data was collected by a Gatan Orius SC1000 CCD camera Model 831 (Gatan Inc., Pleasanton, CA, USA). Image analysis was done using Image J.

### Western blotting and densitometry analyses

SDS-PAGE was run using NuPAGE 4-12% Bis-Tris protein gels with MOPS-SDS buffer (Thermo Fisher Scientific, USA). After electrophoresis, proteins were transferred from gels to Immobilon-P polyvinylidene difluoride 0.45 μm membranes (Millipore, USA) according to manufacturer’s protocol. Membranes were blocked in either Odyssey or Intercept blocking buffer (LI-COR, USA). α-FLAG blots of membrane proteins were probed using α-FLAG antibody (Abcam 2493) as the primary. α-His blots were probed with 6xHis-antibody (Abcam, ab1187) as the primary. For a-glycan blots, hR6 serum from rabbit that binds to the native *C. jejuni* glycan was used as the primary probe (20). A fluorescent goat a-Rabbit IgG IRDye 680RD (LI-COR, USA) was used as the secondary for all blots. Blots were imaged using a LI-COR Odyssey Fc (LI-COR Biosciences, USA). Densitometry was preformed using Image Studio Lite software to measure band intensity. Fluorescence background was subtracted from blots before determining band intensities. For determining membrane protein enrichment (S12/S30), band intensities of membrane proteins for three independent S12 extract replicates and three independent S30 replicates were measured for each protein. The rounded averages of triplicate ratios (S12/S30) and associated error are reported as enrichment in **Fig. 3.** For determining glycoprotein yields from CFGpS reactions, band intensities for glycosylated and aglycosylated bands were obtained from independent, triplicate reactions. The fraction of glycosylated protein and associated standard deviations were calculated via band intensities. To obtain glycoprotein yields, the average fraction glycosylated was multiplied by average total protein yield as calculated from sfGFP fluorescence described below.

### Lipid dye staining and fluorescence immunostaining of vesicles

All reagents used for immunostaining and SEC were sterile filtered with a 0.1 μm filter (Millex-VV Syringe Filter, Merck Millipore Ltd. or Rapid-Flow Filter, Nalgene). To determine vesicle elution fractions, extract was probed with FM 4-64 lipid dye (Life Technologies), a lipophilic styrene dye that has low fluorescence in aqueous solution and becomes brightly fluorescent upon incorporation into membranes. FM-464 dye preferentially stains the inner membrane of *E. coli,* but has been used to dye the outer membrane as well (75, 76). FM 4-64 lipid dye was prepared in stock solutions at 10 mg/mL in 100% DMSO, then diluted 1,000-fold in nuclease free water before use. 80 μL of extract, 10 μL 10x PBS, and 10 μL of FM 4-64 were mixed to a final concentration of 1 ng dye/μL. Samples were incubated with dye in the dark for 10 mins at 37 °C prior to SEC. To verify the presence of glycosylation components in vesicles, we probed for the LLO with a red fluorescent soybean agglutinin (SBA) lectin, a protein complex which specifically binds to the *C. jejuni* LLO (66), and for PglB with an orthogonal green fluorescent α-FLAG antibody as described above. For a-FLAG immunostaining and SBA staining, 90 μL extract and 10 μL of 10xPBS were mixed with 2 μL of a-FLAG-DyLight 488 (Invitrogen, MA191878D488) and 4 μL of SBA-AlexaFluor™ 594 (Invitrogen, 32462). Antibody and SBA were incubated with extract in the dark with agitation overnight at 4 °C prior to SEC.

### Size exclusion chromatography (SEC) of vesicles

100 μL of extract mixture (stained with lipid dye or antibody) was flowed over a size exclusion chromatography column with PBS. Elution fractions were collected into a clear polystyrene 96-well plate (Costar 3370, Corning Inc., USA) at a rate of 0.4 min/well using a Gilson FC 204 Fraction Collector (Gilson, Inc., USA). Poly-Prep chromatography columns (Bio-Rad, USA) were packed with 8 mL of Sepharose 4B resin 45-165 μm bead diameter, (Sigma Aldrich, USA) and washed with sterile PBS 3 times before use. Elution fluorescence was measured using a Synergy H1 microplate reader (Biotek, USA). Excitation and emission wavelengths for SBA-AlexaFluor™ 594 were 590 and 617 nm, respectively. Excitation and emission wavelengths for a-FLAG-DyLight 488 were 493 and 528 nm, respectively. Vesicles stained with FM 4-64 lipid dye were used to determine the characteristic vesicle elution fraction. Reference samples probed with FM 4-64 were used to determine the characteristic vesicle elution fraction in each experiment. For plots, each curve was background subtracted and normalized to the highest RFUs measured for each respective fluorescent elution profile.

### CFE reactions

Protein synthesis was carried out with a modified PANOx-SP system in triplicate reactions, with each reaction containing a uniquely-prepared extract (70). Specifically, 1.5 mL microcentrifuge tubes (Axygen, MCT-150-C) were charged with 15 μL reactions containing 200 ng pJL1-sfGFP plasmid (**SI Appendix, Table S1**), 30% (v/v%) extract and the following: 6 mM magnesium glutamate (Sigma, 49605), 10 mM ammonium glutamate (MP, 02180595), 130 mM potassium glutamate (Sigma, G1501), 1.2 mM adenosine triphosphate (Sigma A2383), 0.85 mM guanosine triphosphate (Sigma, G8877), 0.85 mM uridine triphosphate (Sigma U6625), 0.85 mM cytidine triphosphate (Sigma, C1506), 0.034 mg/mL folinic acid, 0.171 mg/mL *E. coli* tRNA (Roche 10108294001), 2 mM each of 20 amino acids, 30 mM phosphoenolpyruvate (PEP, Roche 10108294001), 0.4 mM nicotinamide adenine dinucleotide (Sigma N8535-15VL), 0.27 mM coenzyme-A (Sigma C3144), 4 mM oxalic acid (Sigma, PO963), 1 mM putrescine (Sigma, P5780), 1.5 mM spermidine (Sigma, S2626), and 57 mM HEPES (Sigma, H3375). To gauge extract CFE productivity, reactions were carried out for 20 hours at 30 °C.

### GFP fluorescence assay

The activity of cell-free-derived sfGFP was determined using an in-extract fluorescence analysis as described previously (39). Briefly, 2 μL of cell-free reaction product was diluted into 48 μL of Ambion nanopure water (Invitrogen, USA). The solution was then placed in a Costar 96-well black assay plate (Corning, USA). Fluorescence was measured using a Synergy H1 microplate reader (Biotek, USA). Excitation and emission wavelengths for sfGFP fluorescence were 485 and 528 nm, respectively.

### Cell-free glycoprotein synthesis (CFGpS) reactions

For crude extract-based expression of glycoproteins, a two-phase scheme was implemented as previously described (34). In this work, protein synthesis was carried out as described above at 15 μL in PCR strip tubes (Thermo Scientific AB-2000) with 50 ng template DNA. Reactions were supplemented with the plasmids encoding permissible or non-permissible sequons on sfGFP acceptor proteins. pJL1-sfGFP-DQNAT-His (permissible) and pJL1-sfGFP-AQNAT-His (non-permissible) were used for PglB-mediated glycosylation; pJL1-sfGFP-MOOR-His (permissible) and pJL1-sfGFP-MOOR_mut_-His (non-permissible) were used for PglO-mediated glycosylation (**SI Appendix, Table S2**). Reactions were set up in triplicate on ice, with each reaction containing a uniquely-prepared extract. CFPS time was measured as the time at which reactions were moved to 30 °C to the time when reactions were spiked with MnCl_2_. In the second phase, protein glycosylation was initiated by the addition of MnCl_2_ at a final concentration of 25 mM. In addition to MnCl_2_ (Sigma 63535), either 0.1% (w/v%) DDM (Anatrace, D310S) or 100 mM sucrose was supplemented to PglB or PglO reactions, respectively. Glycosylation proceeded at 30 °C for 16 hrs. After glycosylation, GFP fluorescence was used to quantitate the total amount of acceptor protein synthesized, and Western blots were used to calculate the fraction of glycosylated and aglycosylated proteins.

### Estimation of vesicle membrane area

The equation below was used to calculate vesicle surface area (m^2^/mL), where Rave is average vesicle radius (m), C is concentration of particles measured by NTA (particles/mL).

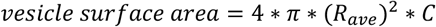

## Acknowledgments

We acknowledge Jessica Stark and Weston Kightlinger for helpful discussions about CFGpS, Aravind Natarajan and Dominic Mills for helpful discussions about *O*-OSTs and their substrate specificities, Ashty Karim for helpful discussions on data visualization, and Han Teng Wong and Charlotte Abrahamson for fruitful discussions regarding immunostaining. We thank Markus Aebi for providing the hR6 serum and Jeff Tabor for providing the DNA encoding NarX. This work made use of the BioCryo facility of Northwestern University’s NUANCE Center, which has received support from the SHyNE Resource (NSF ECCS-1542205), the IIN, and Northwestern’s MRSEC program (NSF DMR-1720139). We gratefully acknowledge support from the Defense Threat Reduction Agency Grant HDTRA1-15-10052/P00001, the National Science Foundation Grants 1936789 and 1844336, the Air Force Research Laboratory Center of Excellence Grant FA8650-15-2-5518, the David and Lucile Packard Foundation, and the Camille Dreyfus Teacher-Scholar Program. This project was also supported in part by fellowships awarded to J.M.H. (NDSEG-36373) and K.F.W. (ND-CEN-013-096) through the National Defense Science and Engineering (NDSEG) Fellowship Program, sponsored by the Air Force Research Laboratory, the Office of Naval Research, and the Army Research Office. J.M.H thanks the Ryan Fellowship awarded by Northwestern University. J.A.P. was supported by an NSF Graduate Research Fellowship. The U.S. Government is authorized to reproduce and distribute reprints for Governmental purposes notwithstanding any copyright notation thereon. The views and conclusions contained herein are those of the authors and should not be interpreted as necessarily representing the official policies or endorsements, either expressed or implied, of the Defense Threat Reduction Agency, or the U.S. Government.

