## Supplemental Information for "Improving cell-free glycoprotein synthesis by characterizing and enriching native membrane vesicles"

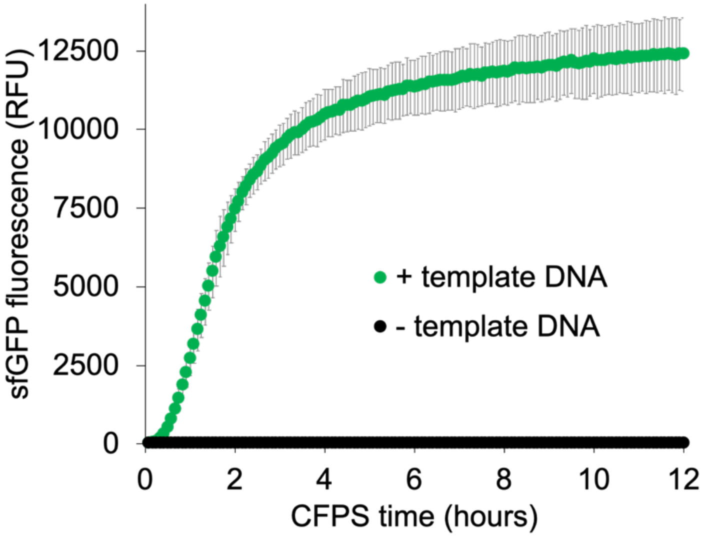


**Figure S1**. **Production of superfolder green fluorescent protein (sfGFP) in standard cell-free expression (CFE) reaction conditions.** Protein synthesis proceeds when template DNA is present, and no fluorescence is observed when template DNA is omitted. Error bars represent standard deviations of three independent CFE reactions.

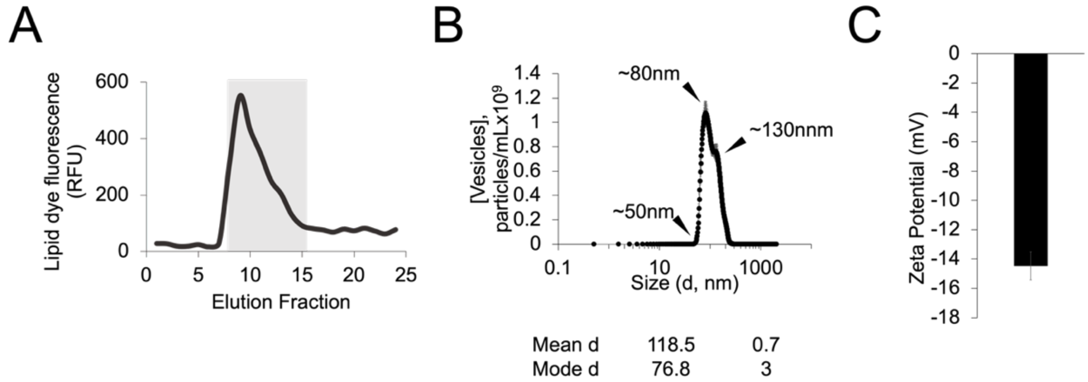


**Figure S2**. **Purification and characterization of membrane vesicles from CFE extracts**. **(A)** SEC chromatogram of extracts probed with FM 4-64 lipid dye. The gray segment indicates the characteristic vesicle elution fraction. **(B)** NTA analysis of purified vesicles collected from fractions 9 and 10 eluted in **(A). (C)** Zeta potential analysis of purified vesicles in PBS. Error bars represent standard deviation of triplicate measurements of purified vesicles.

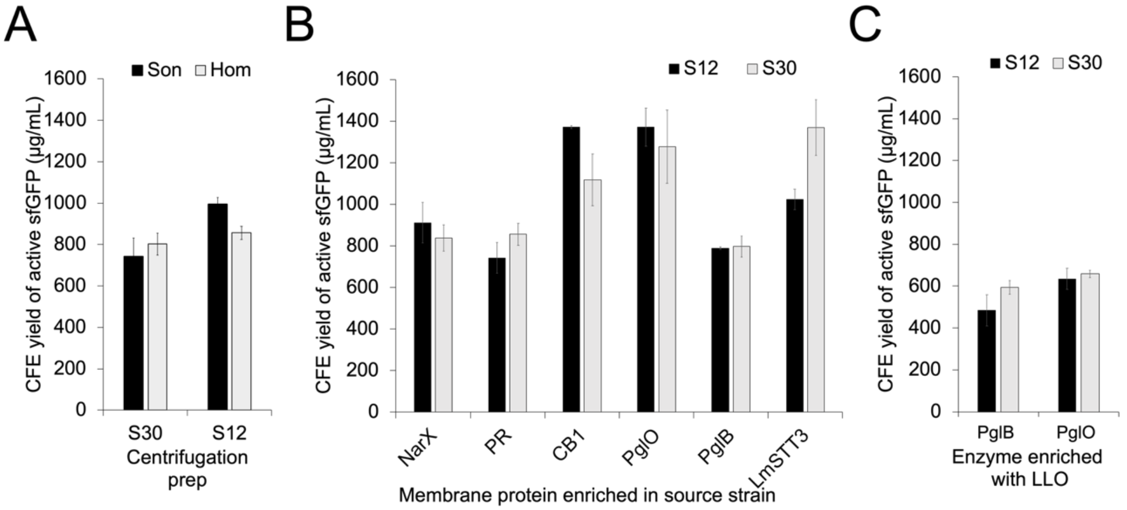


**Figure S3. CFE productivities of all S30 and S12 extracts characterized in this study.** Reactions were run for 20 hours at 30 °C under standard conditions. **(A)** Extracts made from 'blank' chassis strains with no overexpressed components. These extracts are characterized in **Figures 1** and **2** of the main text**. (B)** Extracts enriched with membrane proteins, characterized in **Figure 3** of the main text**. (C)** CFGpS extracts with enrichment of *C. jejuni* LLO and the denoted enzyme in the strain. CFGpS extracts are characterized in **Figure 4** of the main text. Error bars represent standard deviations of triplicate independent reactions.

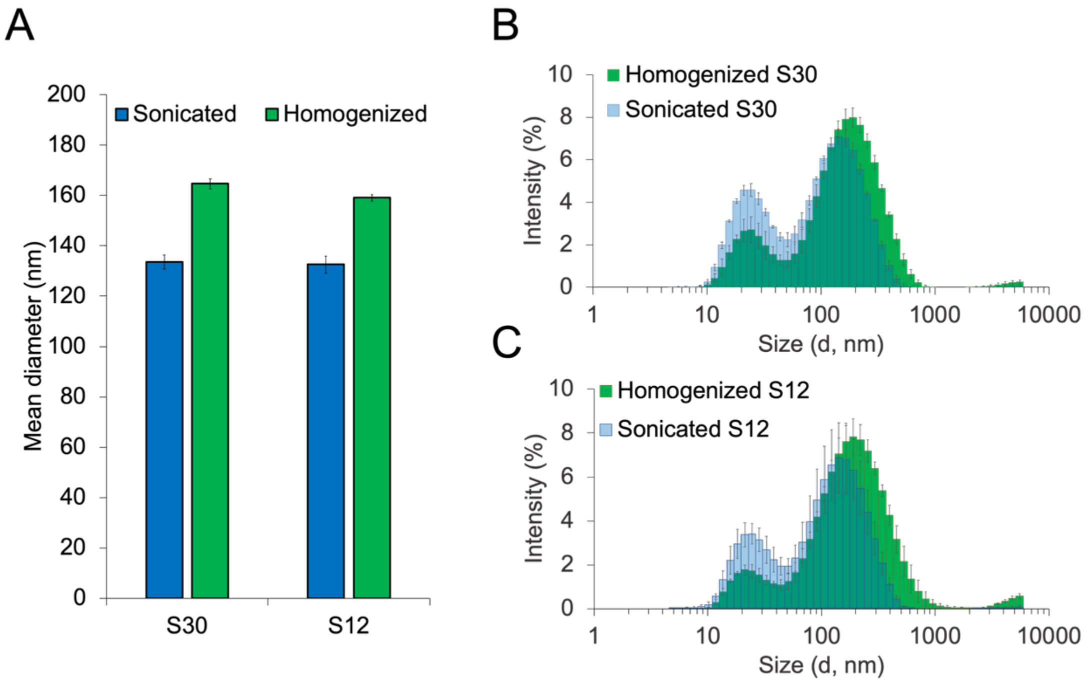


**Figure S4**. **Additional light scattering characterization of extracts presented in Figure 2 of the main text. (A)** Mean vesicle diameters, determined by NTA. DLS analysis of **(B)** S30 and **(C)** S12 extracts. Spectra corroborate larger, right-shifted vesicle peaks in homogenized extracts in both cases. Consistent with NTA particle counting, the relative peak heights of ~20 nm peak (ribosomes/small cellular complexes) to vesicle peak indicates that homogenized extracts contain higher concentrations of vesicles than sonicated extracts for each given prep method. Error bars represent the standard deviation of measurements of three independent extracts.

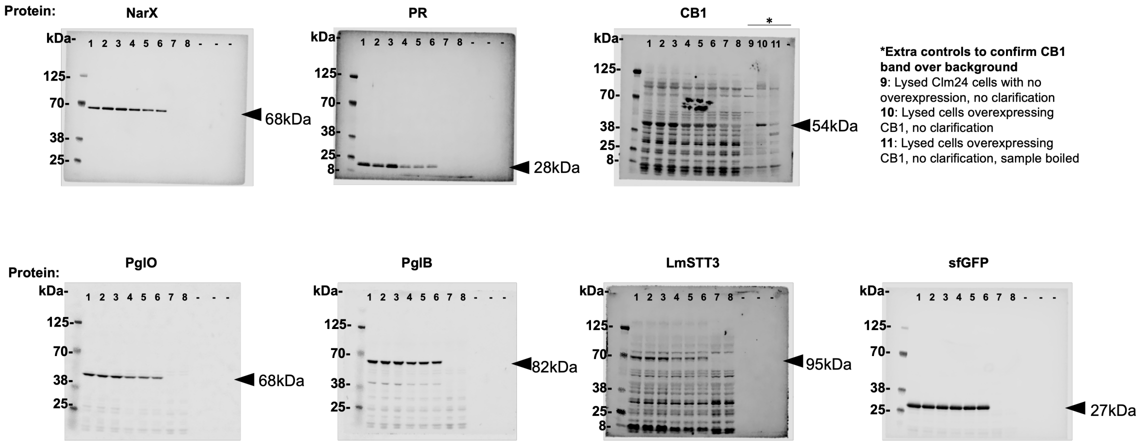


| Lane | S12/S30 | Extract replicate (#) | Notes |
|---|---|---|---|
| Ladder | - | - | - |
| 1 | S12 | 1 | - |
| 2 | S12 | 2 | - |
| 3 | S12 | 3 | - |
| 4 | S30 | 1 | - |
| 5 | S30 | 2 | - |
| 6 | S30 | 3 | - |
| 7 | S12 | N/A | No overexpression, strain background control |
| 8 | S30 | N/A | No overexpression, strain background control |

**Figure S5**. **Western blot analysis of membrane protein enrichment in S30 and S12 extracts.** Uncropped α-FLAG blots against each of the indicated recombinant proteins. The theoretical mass of each recombinant protein is listed next to black arrows, indicating the corresponding band. We observe the well-documented effect that membrane proteins run anomalously on SDS-PAGE, running 'light' with respect to the protein ladder standard. A lane key is presented below. Note that extra controls (indicated to the right of blot) were needed for the CB1 blot to confirm the presence of the protein.

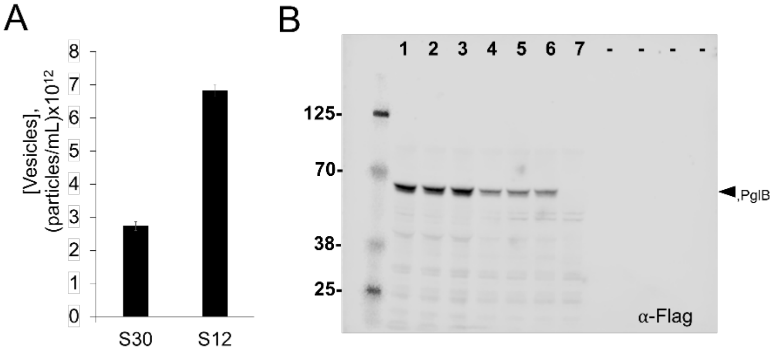


| Lane | S12/S30 | Extract replicate (#) | Notes |
|---|---|---|---|
| Ladder | - | - | - |
| 1 | S12 | 1 | - |
| 2 | S12 | 2 | - |
| 3 | S12 | 3 | - |
| 4 | S30 | 1 | - |
| 5 | S30 | 2 | - |
| 6 | S30 | 3 | - |
| 7 | S12 | N/A | LLO only expression control |

**Figure S6**. **Characterization of vesicle and PglB enrichment in CFGpS extracts. (A)** Concentrations of vesicles in extracts enriched with PglB and *C. jejuni* LLO, measured via NTA. Error bars represent standard deviation of measurements of three independent extracts. **(B)** α-FLAG Western blot against PglB in an extract enriched with both PglB and LLO. Corresponding lane key is to the right of the gel.

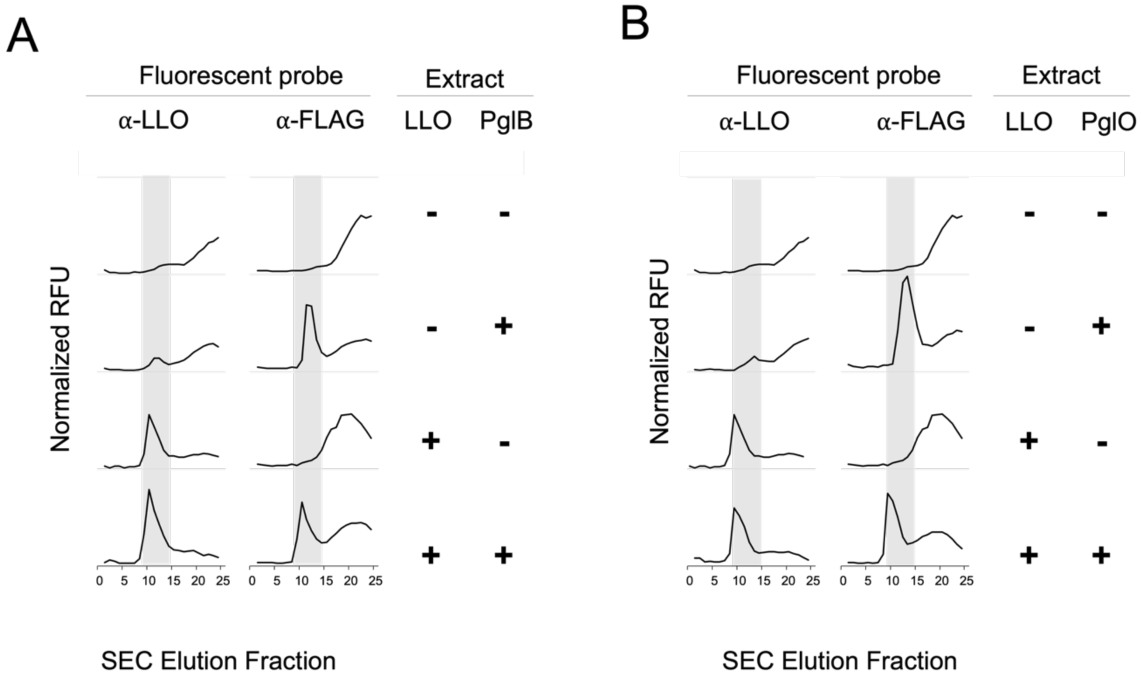


**Figure S7**. **Fluorescent probes confirm that glycosylation components are embedded in membrane vesicles in CFE extracts**. Fluorescence SEC chromatograms of vesicles probed with α-LLO and α-OST reagents are presented. Vesicle elution fraction is highlighted in gray. Analysis of extracts and controls with the *N*-linked PglB OST are presented in **(A)** and for the *O*-linked PglO OST in **(B)**.

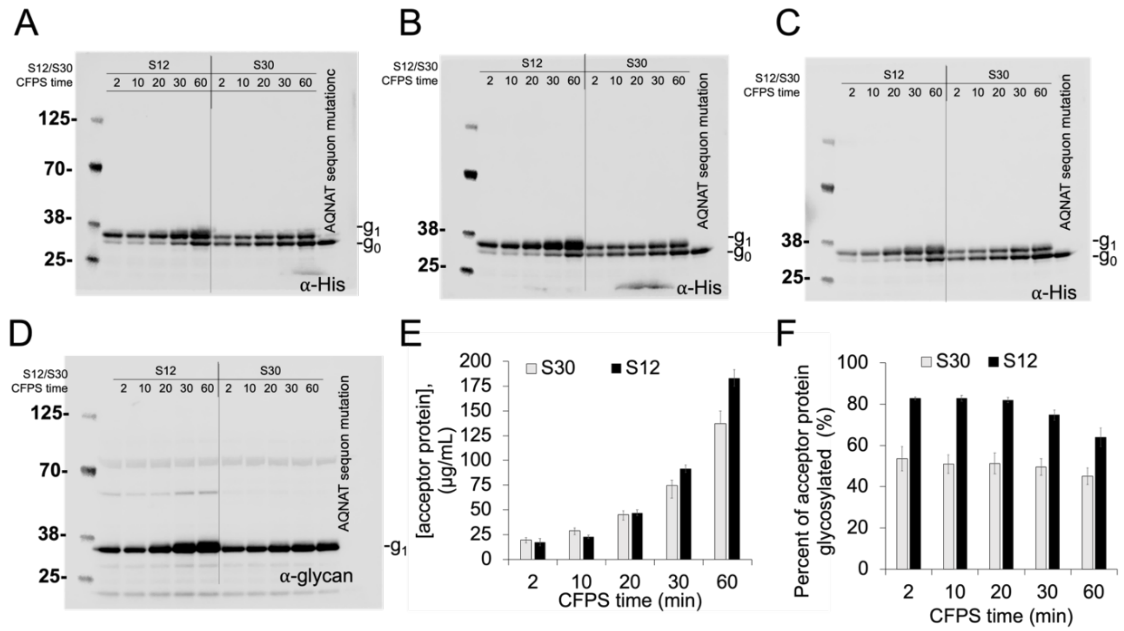


**Figure S8**. **Characterization of *N*-linked glycosylation in S12 and S30 CFGpS extracts.** Triplicate α-His Western blots against CFGpS-derived acceptor proteins are shown in **(A)-(C)** corresponding with extract replicates 1-3. Western blots in **(A)-(C)** were used to calculate glycoprotein yields in **Figure 4** of the main text. **(D)** α-glycan blot of the corresponding reactions in **(A).** $g_o$ denotes aglycosylated acceptor protein and $g_1$ indicates glycoprotein. **(E)** Total acceptor protein produced, and **(F)** percent of acceptor protein converted to glycoprotein at each condition. Error bars represent standard deviation of three independent reactions.

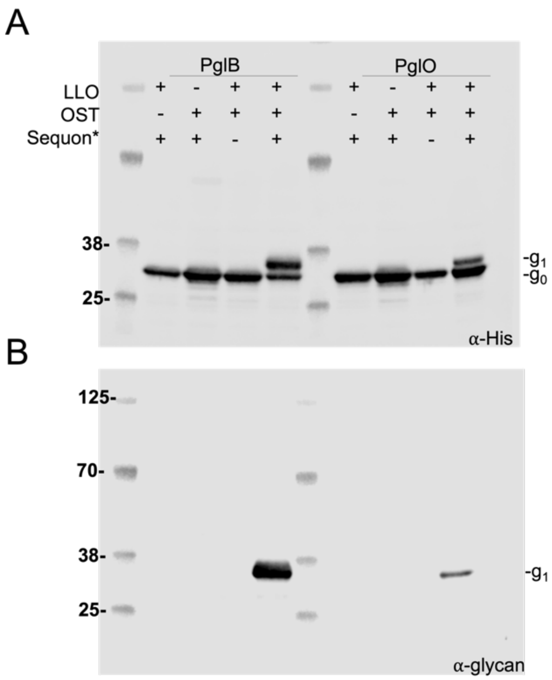


*PglB permissible sequon: DQNAT
PglB impermissible sequon: AQNAT
PglO permissible sequon: WPAAASAP
PglO impermissible sequon: WPAAAGAP

**Figure S9. Residue-specific glycosylation of acceptor proteins with permissible sequons for PglB and PglO.** Generally, glycosylation preferences of PglO are less understood then those for PglB. PglB samples are used as blotting references for positive and negative controls, as the sequon specificities for positive and negative sequons are well-characterized. **(A)** α-His blot of CFGpS reactions run with a 5 minute CFPS time. Glycosylated band is only present when all glycosylation components and a permissible sequon are present. $g_o$ denotes aglycosylated acceptor protein and $g_1$ indicates glycoprotein. **(B)** α-glycan blot that corresponds to data from (**A**), showing a signal only for $g_1$ when all glycosylation components are present.

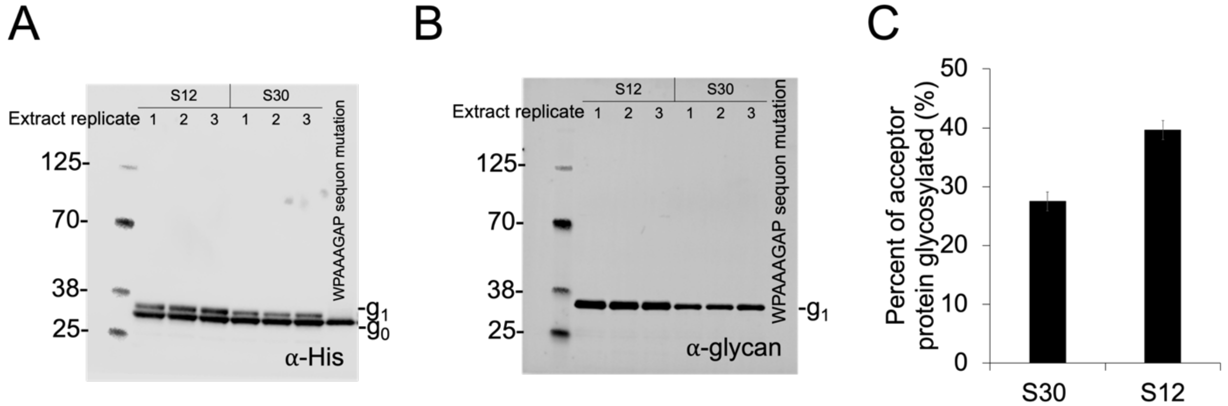


**Figure S10. Characterization of *O*-linked glycosylation in CFGpS extracts**. **(A)** α-His Western blot of PglO CFGpS reactions run with 20 minute CFPS times and **(B)** α-glycan blot corresponding to data in **(A)**. $g_0$ denotes aglycosylated acceptor protein and $g_1$ indicates glycoprotein. **(C)** Percent of acceptor protein converted to glycoprotein. Error bars represent standard deviation of three independent CFGpS reactions.

**Table S1.** Information on proteins selected for extract enrichment in this study.

| Protein | Taxonomical origin | Predicted # TM helices | Function(s) | Size (kDa) | UniProt ID |
|---|---|---|---|---|---|
| PglB | *Campylobacter jejuni* | 13 | Catalyzes *N*-linked glycosylation | 82 | Q5HTX9 |
| PglO | *Neisseria gonorrhoeae* | 11 | Catalyzes *O*-linked glycosylation | 68 | Q5FA54 |
| NarX | *Escherichia coli* | 2 | Signal transduction for nitrate biosensing | 68 | P0AFA2 |
| Proteorhodopsin (PR) | Uncultured marine gamma proteobacterium EBAC3108 | 7 | Green light absorbing proteorhodopsin | 28 | Q9F7P4 |
| Cannabinoid receptor 1 (CB1) | *Homo sapiens* | 7 | G protein coupled receptor, molecular sensing | 54 | P21554 |
| STT3D | *Leishmania major* | 11 | Catalyzes *N*-linked glycosylation | 95 | E9AET9 |

**Table S2**. Acceptor protein coding sequences used in CFGpS reactions.

| Sequon | Coding Sequence |
|---|---|
| pJL1-sfGFP-DQNAT | ATGAGCAAAGGTGAAGAACTGTTTACCGGCGTTGTGCCGATTCTGGTGGAACTGGATGGCGAT<br>GTGAACGGTCACAAATTCAGCGTGCGTGGTGAAGGTGAAGGCGATGCCACGATTGGCAAACT<br>GACGCTGAAATTTATCTGCACCACCGGCAAACTGCCGGTGCCGTGGCCGACGCTGGTGACCA<br>CCCTGACCTATGGCGTTCAGTGTTTTAGTCGCTATCCGGATCACATGAAACGTCACGATTTCTT<br>TAAATCTGCAATGCCGGAAGGCTATGTGCAGGAACGTACGATTAGCTTTAAAGATGATGGCAAA<br>TATAAAACGCGCGCCGTTGTGAAATTTGAAGGCGATACCCTGGTGAACCGCATTGAACTGAAA<br>GGCACGGATTTTAAAGAAGATGGCAATATCCTGGGCCATAAACTGGAATACAACTTTAATAGCC<br>ATAATGTTTATATTACGGCGGATAAACAGAAAAATGGCATCAAAGCGCAGTTTACCGTTCGCCA<br>TAACGTTGAAGATGGCAGTGTGCAGCTGGCAGATCATTATCAGCAGAATACCCCGATTGGTGA<br>TGGTCCGGTGCTGCTGCCGGATAATCATTATCTGAGCACGCAGACCGTTCTGTCTAAAGATCC<br>GAACGAAAAAGGCACGCGGGACCACATGGTTCTGCACGAATATGTGAATGCGGCAGGTATTAC<br>GCTAGGTGCGGCCGCAGAACAAAAACTCATCTCAGAAGAGGATCTGAATGGGGCCGCACTCG<br>AGGGTGGCGATCAGAACGCGACCGGCGGTCATCACCATCATCACCATTAA |
| pJL1-sfGFP AQNAT | ATGAGCAAAGGTGAAGAACTGTTTACCGGCGTTGTGCCGATTCTGGTGGAACTGGATGGCGAT<br>GTGAACGGTCACAAATTCAGCGTGCGTGGTGAAGGTGAAGGCGATGCCACGATTGGCAAACT<br>GACGCTGAAATTTATCTGCACCACCGGCAAACTGCCGGTGCCGTGGCCGACGCTGGTGACCA<br>CCCTGACCTATGGCGTTCAGTGTTTTAGTCGCTATCCGGATCACATGAAACGTCACGATTTCTT<br>TAAATCTGCAATGCCGGAAGGCTATGTGCAGGAACGTACGATTAGCTTTAAAGATGATGGCAAA<br>TATAAAACGCGCGCCGTTGTGAAATTTGAAGGCGATACCCTGGTGAACCGCATTGAACTGAAA<br>GGCACGGATTTTAAAGAAGATGGCAATATCCTGGGCCATAAACTGGAATACAACTTTAATAGCC<br>ATAATGTTTATATTACGGCGGATAAACAGAAAAATGGCATCAAAGCGCAGTTTACCGTTCGCCA<br>TAACGTTGAAGATGGCAGTGTGCAGCTGGCAGATCATTATCAGCAGAATACCCCGATTGGTGA<br>TGGTCCGGTGCTGCTGCCGGATAATCATTATCTGAGCACGCAGACCGTTCTGTCTAAAGATCC<br>GAACGAAAAAGGCACGCGGGACCACATGGTTCTGCACGAATATGTGAATGCGGCAGGTATTAC<br>GCTAGGTGCGGCCGCAGAACAAAAACTCATCTCAGAAGAGGATCTGAATGGGGCCGCACTCG<br>AGGGTGGCGCTCAGAACGCGACCGGCGGTCATCACCATCATCACCATTAA |
| pJL1-sfGFP-MOOR | ATGAGCAAAGGTGAAGAACTGTTTACCGGCGTTGTGCCGATTCTGGTGGAACTGGATGGCGAT<br>GTGAACGGTCACAAATTCAGCGTGCGTGGTGAAGGTGAAGGCGATGCCACGATTGGCAAACT<br>GACGCTGAAATTTATCTGCACCACCGGCAAACTGCCGGTGCCGTGGCCGACGCTGGTGACCA<br>CCCTGACCTATGGCGTTCAGTGTTTTAGTCGCTATCCGGATCACATGAAACGTCACGATTTCTT<br>TAAATCTGCAATGCCGGAAGGCTATGTGCAGGAACGTACGATTAGCTTTAAAGATGATGGCAAA<br>TATAAAACGCGCGCCGTTGTGAAATTTGAAGGCGATACCCTGGTGAACCGCATTGAACTGAAA<br>GGCACGGATTTTAAAGAAGATGGCAATATCCTGGGCCATAAACTGGAATACAACTTTAATAGCC<br>ATAATGTTTATATTACGGCGGATAAACAGAAAAATGGCATCAAAGCGAATTTTACCGTTCGCCAT<br>AACGTTGAAGATGGCAGTGTGCAGCTGGCAGATCATTATCAGCAGAATACCCCGATTGGTGAT<br>GGTCCGGTGCTGCTGCCGGATAATCATTATCTGAGCACGCAGACCGTTCTGTCTAAAGATCCG<br>AACGAAAAAGGCACGCGGGACCACATGGTTCTGCACGAATATGTGAATGCGGCAGGTATTACG<br>GGCTCTTCTGGAGGGTCTGGCGATCCACGCAATGTGGGTGGGGATTTGGACTGGCCGGCGG<br>CAGCGAGTGCACCTCAACCCGGTAAACCTCCTCGTCATCACCACCATCATCACTAA |
| pJL1-sfGFP-MOORmut | ATGAGCAAAGGTGAAGAACTGTTTACCGGCGTTGTGCCGATTCTGGTGGAACTGGATGGCGAT<br>GTGAACGGTCACAAATTCAGCGTGCGTGGTGAAGGTGAAGGCGATGCCACGATTGGCAAACT<br>GACGCTGAAATTTATCTGCACCACCGGCAAACTGCCGGTGCCGTGGCCGACGCTGGTGACCA<br>CCCTGACCTATGGCGTTCAGTGTTTTAGTCGCTATCCGGATCACATGAAACGTCACGATTTCTT<br>TAAATCTGCAATGCCGGAAGGCTATGTGCAGGAACGTACGATTAGCTTTAAAGATGATGGCAAA<br>TATAAAACGCGCGCCGTTGTGAAATTTGAAGGCGATACCCTGGTGAACCGCATTGAACTGAAA<br>GGCACGGATTTTAAAGAAGATGGCAATATCCTGGGCCATAAACTGGAATACAACTTTAATAGCC<br>ATAATGTTTATATTACGGCGGATAAACAGAAAAATGGCATCAAAGCGAATTTTACCGTTCGCCAT<br>AACGTTGAAGATGGCAGTGTGCAGCTGGCAGATCATTATCAGCAGAATACCCCGATTGGTGAT<br>GGTCCGGTGCTGCTGCCGGATAATCATTATCTGAGCACGCAGACCGTTCTGTCTAAAGATCCG<br>AACGAAAAAGGCACGCGGGACCACATGGTTCTGCACGAATATGTGAATGCGGCAGGTATTACG<br>GGCTCTTCTGGAGGGTCTGGCGATCCACGCAATGTGGGTGGGGATTTGGACTGGCCGGCGG<br>CAGCGGGTGCACCTCAACCCGGTAAACCTCCTCGTCATCACCACCATCATCACTAA |

**Table S3.** Strains and plasmids used in this study.

| Strain/Plasmid | Description | Antibiotic Resistance | Reference |
|---|---|---|---|
| *E. coli* CLM24 | W3110, ΔWaaL | N/A | (1) |
| pJL1-sfGFP | pJL1 plasmid encoding superfolder GFP | Kan50 | (2) |
| pSF-CjPglB- | pSN18 derivative encoding *C. jejuni* PglB with C-terminal FLAG epitope tag | Carb100 | (3) |
| pSF-NgPglO | pSN18 derivative encoding *N. gonorrhoeae* PglO with C-terminal FLAG epitope tag | Carb100 | This study |
| pSF-EcNarX | pSN18 derivative encoding *E. coli* NarX with C-terminal FLAG epitope tag | Carb100 | This study |
| pSF-PR | pSN18 derivative encoding Uncultured marine gamma proteobacterium EBAC3108 proteorhodopsin with C-terminal FLAG epitope tag | Carb100 | This study |
| pSF-HsCB1 | pSN18 derivative encoding *H. sapiens* Cannabinoid receptor 1 with C-terminal FLAG epitope tag | Carb100 | This study |
| pSF-LmSTT3D | pSN18 derivative encoding *L. major* STT3D with C-terminal FLAG epitope tag | Carb100 | This study |
| pSF-sfGFP | pSN18 derivative encoding superfolder GFP with C-terminal FLAG epitope tag | Carb100 | This study |
| pMW07-pglΔB | pMW07 plasmid encoding *C. jejuni* protein glycosylation locus (pgl) with complete in-frame deletion of CjPglB | Cm34 | (3) |
| pJL1-sfGFP-DQNAT | pJL1 plasmid encoding superfolder GFP modified at the C-terminus with 30 amino acids containing an optimal DQNAT sequon followed by a 6x-His tag | Kan50 | This study, see **Table S2** for sequence |
| pJL1-sfGFP-AQNAT | pJL1 plasmid encoding superfolder GFP modified at the C-terminus with 30 amino acids containing a non-permissible AQNAT sequon followed by a 6x-His tag | Kan50 | This study, see **Table S2** for sequence |
| pJL1-sfGFP-MOOR | pJL1 plasmid encoding superfolder GFP modified at the C-terminus with 32 amino acids containing the minimum optimal *O*-linked recognition site (MOOR)[4] followed by a 6x-His tag | Kan50 | This study, see **Table S2** for sequence |
| pJL1-sfGFP-MOORmut | pJL1 plasmid encoding superfolder GFP modified at the C-terminus with 32 amino acids containing a non-permissible minimum optimal *O*-linked recognition site (MOORmut) followed by a 6x-His tag | Kan50 | This study, see **Table S2** for sequence |

### SI Methods

#### *Zeta Potential Analysis*

Zeta potential measurements were performed in triplicate for 15 scans per measurement on a Zetasizer Nano ZS (Malvern Instruments Ltd.) using standard settings at room temperature and in disposable zeta potential cuvettes (Malvern Instruments Ltd., UK DTS1070).
